# Transcriptome-wide map of m^6^A circRNAs identified in hypoxic pulmonary hypertension rat model

**DOI:** 10.1101/625178

**Authors:** Hua Su, Lin Zhou, Na Li, Guowen Wang, Lingfang Wu, Xiuqing Ma, Kejing Ying, Ruifeng Zhang

## Abstract

Hypoxic pulmonary hypertension (HPH) is a lethal disease. CircRNAs and m^6^A circRNAs have been reported to be associated with cancer progression, but the expression profiling of m^6^A circRNAs has not been identified in HPH. This study was to investigate the transcriptome-wide map of m^6^A circRNAs in HPH. In this study, hypoxia-induced PH rat model was established. Total RNA was extracted and purified from lungs of rats, then circRNAs were detected and annotated by RNA-seq analysis. m^6^A RNA Immunoprecipitation (MeRIP) was performed following rRNA depletion, and RNA-seq library was constructed. CircRNA–miRNA–mRNA co-expression network was also constructed. In vitro, total m^6^A was measured. m^6^A circXpo6 and m^6^A circTmtc3 were detected in pulmonary artery smooth muscle cells (PASMCs) and pulmonary artery endothelial cells (PAECs) exposed to 21% O_2_ and 1% O_2_ for 48 h, respectively. m^6^A abundance in 166 circRNAs was significantly upregulated and m^6^A abundance in 191 circRNAs was significantly downregulated in lungs of HPH rats. m^6^A abundance in circRNAs was significantly reduced in hypoxia *in vitro*. m^6^A circRNAs were mainly derived from single exons of protein-coding genes. m^6^A influenced the circRNA–miRNA–mRNA co-expression network in hypoxia. m^6^A circXpo6 and m^6^A circTmtc3 were downregulated in hypoxia. In general, our study firstly identified the transcriptome-wide map of m^6^A circRNAs in HPH. m^6^A level in circRNAs was decreased in lungs of HPH rats and in PASMCs and PAECs exposed to hypoxia. Downregulated or upregulated m^6^A level influenced circRNA–miRNA–mRNA co-expression network in HPH. Moreover, we firstly identified two downregulated m^6^A circRNAs in HPH: circXpo6 and circTmtc3. We suggested that m^6^A circRNAs may be used as a potential diagnostic marker or therapy target in the future.

**Author summary:** HPH is a disease with great morbidity and mortality. It is often caused by chronic hypoxic lung diseases, such as chronic obstructive pulmonary disease and interstitial lung diseases. It lacks effective therapy methods so far. CircRNAs are a type of non-coding RNAs and can be used as biomarkers because they are differentially enriched in specific cell types or tissues and not easily degraded. m^6^A is identified as the most universal modification on non-coding RNAs in eukaryotes. CircRNAs can be modified by m^6^A. m^6^A circRNAs in HPH is not well understood yet. Here we identify the transcriptome-wide map of m^6^A circRNAs in HPH. We elucidate that m^6^A level in circRNAs is decreased in lungs of HPH rats and in PASMCs and PAECs exposed to hypoxia. We find that downregulated or upregulated m^6^A level influences circRNA– miRNA–mRNA co-expression network in HPH. Moreover, we are the first to identify two downregulated m^6^A circRNAs in HPH: circXpo6 and circTmtc3. We suggest that m^6^A circRNAs may be used as a potential diagnostic marker or therapy target in the future.

## Introduction

Pulmonary hypertension (PH) is a lethal disease and defined as an increase in the mean pulmonary arterial pressure ≥ 25 mmHg at rest, as measured by right heart catheterization (1). Hypoxic pulmonary hypertension (HPH) belongs to group III PH according to the comprehensive clinical classification of PH, normally accompanied by severe chronic obstructive pulmonary disease (COPD) and interstitial lung diseases (2). HPH is a progressive disease induced by chronic hypoxia (CH) (1). CH triggers over-proliferation of pulmonary artery endothelial cells (PAECs) and pulmonary artery smooth muscle cells (PASMCs), and activation of quiescent fibroblasts, the hallmark of HPH (1, 3). The pathological characteristics of HPH are pulmonary vascular remolding, pulmonary hypertension, and right ventricular hypertrophy (RVH) (4). So far there is no effective therapy for HPH (2). More effective therapeutic targets are needed to be discovered.

Circular RNAs (circRNAs) were firstly found abundant in eukaryotes using RNA-seq approach (5–7). Pre-mRNA is spliced with the 5’ and 3’ ends, forming a ‘head-to-tail’ splice junction, then circRNAs are occurred (5). According to the genome origin, circRNAs may be classified into four different subtypes: exonic circRNA (ecircRNA), intronic circRNA (ciRNA), exon–intron circRNA (EIciRNA) and tRNA introns circRNA (tricRNA) (5). CircRNAs are reported to play crucial roles in miRNA binding, protein binding, regulation of transcription, and post-transcription (5, 8). Recent reports indicated that circRNAs can translate to proteins (8, 9). Moreover, circRNAs are widely expressed in human umbilical venous endothelial cells when stimulated by hypoxia (10, 11). Up to date, only a few reports mentioned PH-associated circRNAs. CircRNAs expression profile is demonstrated in HPH and chronic thromboembolic pulmonary hypertension (12). However, it is still unknown that the post-transcript modification of circRNAs in HPH.

N^6^-methyladenosine (m^6^A) is regarded as one part of “epitranscriptomics’’ and identified as the most universal modification on mRNAs and noncoding RNAs (ncRNAs) in eukaryotes (13, 14). DRm^6^ACH (D denotes A, U or G; R denotes A, G; H denotes A, C, or U) is a consensus motif occurred in m^6^A modified RNAs (15–17). m^6^A modification is mainly enriched around the stop codons, at 3’untranslated regions (3’ UTRs) and within internal long exons (17–19). Several catalyzed molecules act as “writers”, “readers”, and “erasers” to regulate the m^6^A modification status (14). The methyltransferase complex is known as writers, including methyltransferase-like-3, −14 and −16 (METTL3/METTL14/METTL16), Wilms tumour 1-associated protein (WTAP), RNA binding motif protein 15 (RBM15), vir like m^6^A methyltransferase associated (KIAA1429) and zinc finger CCCH-type containing 13 (ZC3H13), appending m^6^A on DRACH (17, 20, 21). METTL3 is regarded as the core catalytically active subunit, while METTL14 and WTAP play a structural role in METTL3’s catalytic activity (18, 22). The “erasers”, fat mass and obesity related protein (FTO) and alkylation repair homolog 5 (ALKBH5), catalyze the N-alkylated nucleic acid bases oxidatively demethylated (22). The “readers”, the YT521-B homology (YTH) domain-containing proteins family includes YTHDF (YTHDF1, YTHDF2, YTHDF3), YTHDC1, and YTHDC2, specifically recognizes m^6^A and regulates splicing, localization, degradation and translation of RNAs (14, 22, 23). The YTHDF1 and YTHDF2 crystal structures forms an aromatic cage to recognize m^6^A sites in cytoplasm (24). YTHDC1 is the nuclear reader and YTHDC2 binds m^6^A under specific circumstances or cell types (24). Hypoxia may alter the balance of writers-erasers-readers and induce tumor growth, angiogenesis, and progression (25, 26).

Interestingly, circRNAs can be m^6^A-modified. m^6^A circRNAs displayed cell-type-specific methylation patterns in human embryonic stem cells (hESCs) and HeLa cells (14). CircRNAs contained m^6^A modifications are likely to promote protein translation in a cap-independent pattern (9). However, m^6^A circRNAs has not been elucidated in HPH yet. Here we are the first to identify the correlation between m^6^A modification and circRNAs abundance in HPH.

## Results

### m^6^A level of circRNAs was reduced in HPH rats and most circRNAs contained one m^6^A peak

3 weeks treatment by hypoxia resulted in right ventricular systolic pressure (RVSP) elevating to 42.23 ± 1.96 mmHg compared with 27.73 ± 1.71 mmHg in the control (p < 0.001, Fig 1A and 1B). RVH was indicated by the increase of the ratio of the right ventricle (RV), left ventricular plus ventricular septum (LV + S) [RV/ (LV + S)] compared with the control (0.25 ± 0.03 vs. 0.44 ± 0.04, p = 0.001, Fig 1C). The medial wall of the pulmonary small arteries was also significantly thickened (19.28 ± 2.19% vs. 39.26 ± 5.83%, p < 0.001, Fig 1D and 1E). Moreover, in the normoxia group, 53.82 ± 3.27% of the arterioles were non-muscularized (NM) vessels, and 25.13 ± 1.83% were fully muscularized (FM) vessels. In contrast, partially muscularized vessels (PM) and FM vessels showed a greater proportion (32.88 ± 3.15% and 41.41 ± 3.35%) in HPH rats, while NM vessels occupied a lower proportion (25.71 ± 2.55%) (Fig 1F). Fig 1G displayed the heatmap of m^6^A circRNAs expression profiling in normoxia (N) and hypoxia (HPH). m^6^A abundance in 166 circRNAs was significantly upregulated. Meanwhile, m^6^A abundance in 191 circRNAs was significantly downregulated (**S1 Table**, filtered by fold change ≥ 4 and p ≤ 0.00001). Lungs of N and HPH rats were selected to measure m^6^A abundance in purified circRNAs. The m^6^A level in total circRNAs isolated from lungs of HPH rats was lower than that from controls (Fig 1H). Moreover, over 50% circRNAs contained only one m^6^A peak either in lungs of N or HPH rats (Fig 1I).

**Fig 1.**
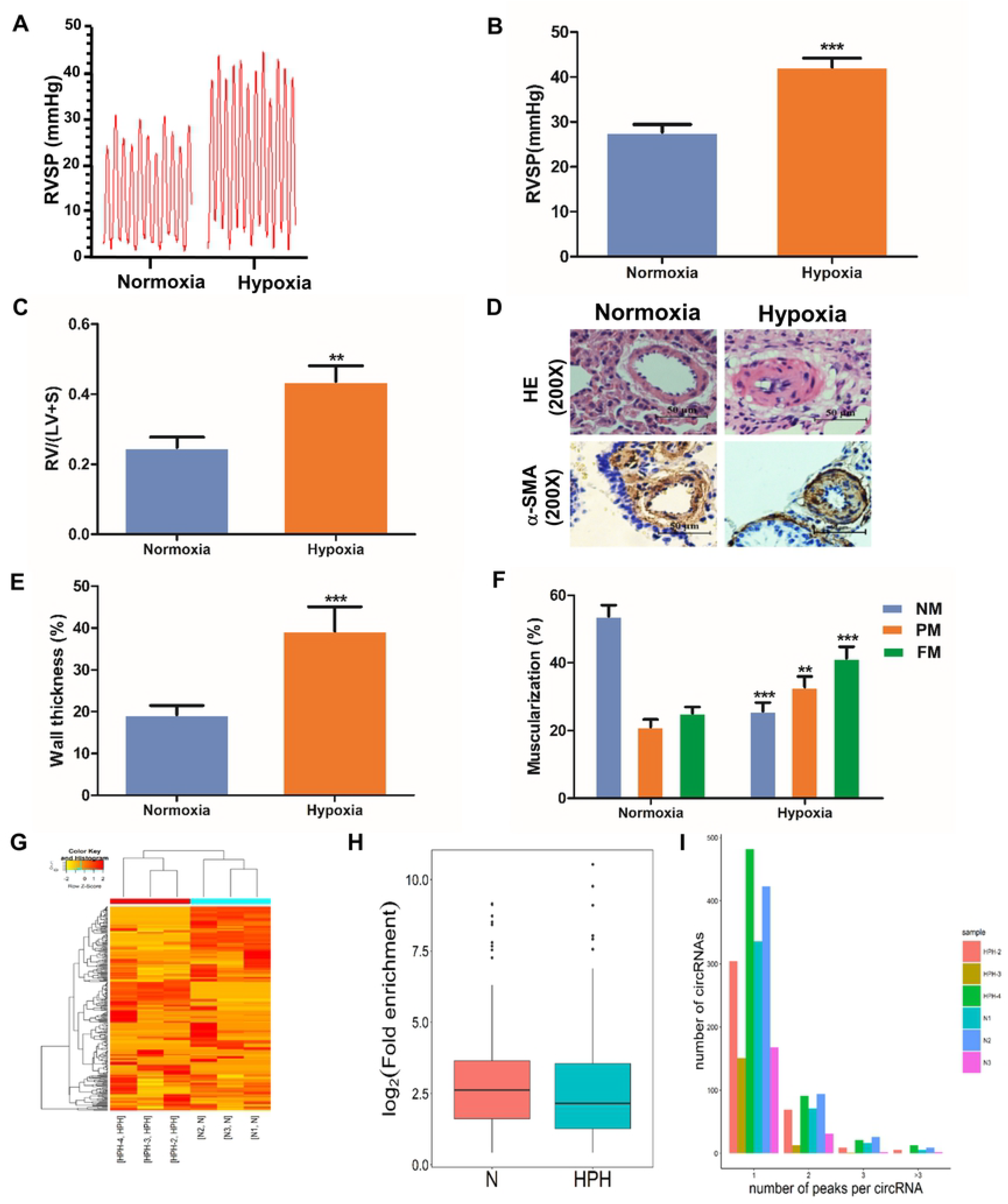
m^6^A level of circRNAs in HPH rats and the number of m^6^A peak in circRNAs. Rats were maintained in a normobaric normoxic (FiO_2_ 21%) or hypoxic (FiO_2_ 10%) chamber for 3 weeks, then RVSP was detected (A, B). (C) The ratio of RV/(LV+S). (D) H&E staining and immunohistochemical staining of α-SMA were performed in the lung sections. Representative images of pulmonary small arteries. Scale bar = 50 μm. Quantification of wall thickness (E) and vessel muscularization (F). (G) Heatmap depicting hierarchical clustering of altered m^6^A circRNAs in lungs of normal (N) and hypoxic pulmonary hypertension (HPH) rats. Red represents higher expression and yellow represents lower expression level. (H) Box-plot for m^6^A peaks enrichment in circRNAs in N and HPH. (I) The distribution of the number of circRNAs (y axis) was plotted based on the number of m^6^A peaks in circRNAs (x axis) in N and HPH. Values are presented as means ± SD (n = 6 in each group). Only vessels with diameter between 30 and 90 μm were analyzed. NM, nonmuscularized vessels; PM, partially muscularized vessels; FM, fully muscularized vessels. **0.001 ≤ p ≤ 0.009 (different from N); ***p < 0.001 (different from N).

### m^6^A circRNAs were mainly from protein-coding genes spanned single exons in N and HPH groups

We analyzed the distribution of the parent genes of total circRNAs, m^6^A-circRNAs, and non-m^6^A circRNAs in N and HPH, respectively. N and HPH groups showed a similar genomic distribution of m^6^A circRNAs and non-m^6^A circRNAs (Fig 2A and 2B). Moreover, about 80% of m^6^A circRNAs and non-m^6^A circRNAs were derived from protein-coding genes in both groups. A previous report indicated that most circRNAs originated from protein-coding genes spanned two or three exons (14). While in our study, over 50% and 40% of total circRNAs from protein-coding genes spanned one exon in N and HPH groups, respectively (Fig 2C and 2D). Similarly, m^6^A circRNAs and non-m^6^A circRNAs were mostly encoded by single exons. Therefore, it was indicated that m^6^A methylation was abundant in circRNAs originated from single exons in N and HPH groups.

**Fig 2.**
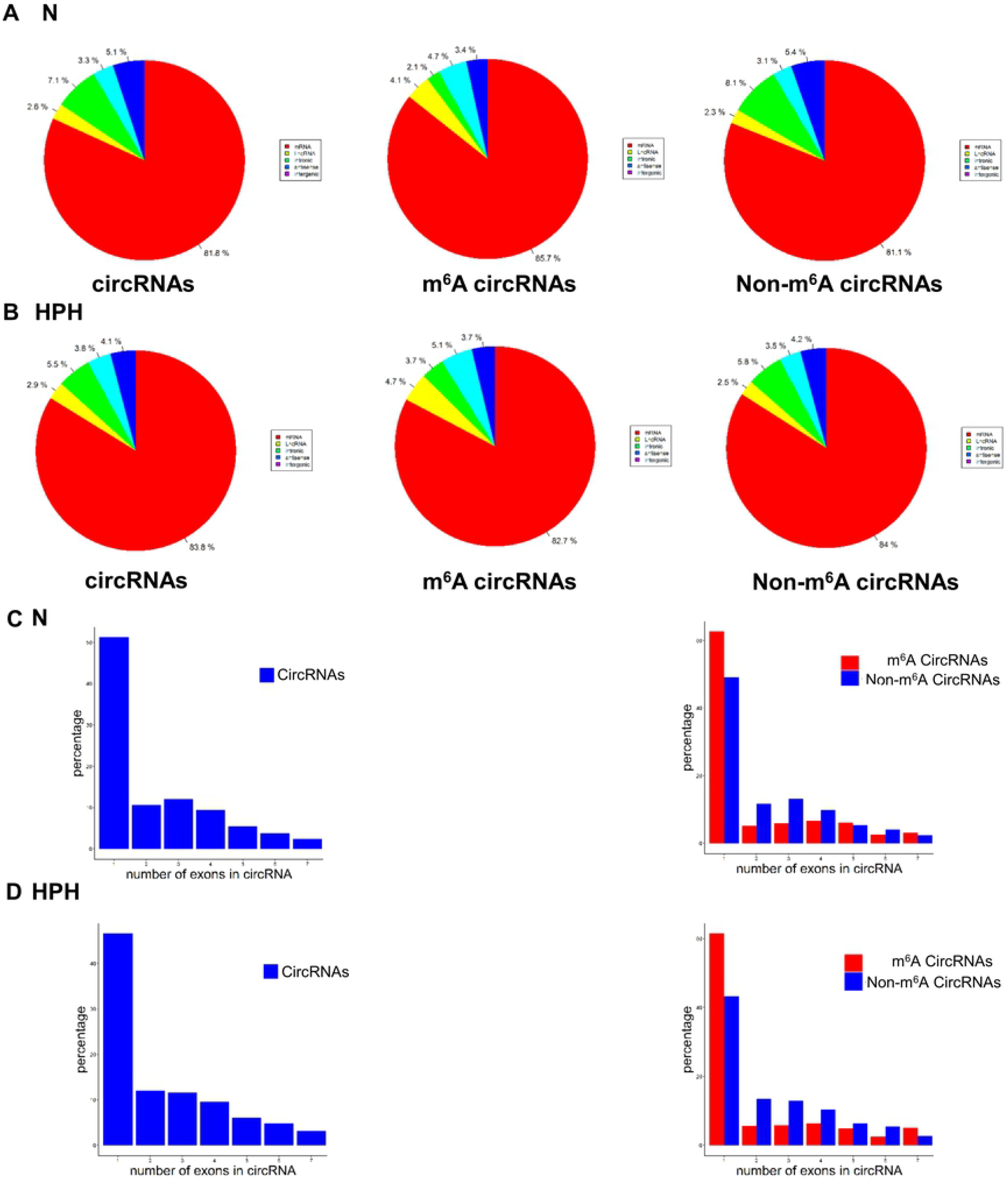
The genomic origins of m^6^A circRNAs. The distribution of genomic origins of total circRNAs (input, left), m^6^A circRNAs (eluate, center), and non-m^6^A circRNAs (supernatant, right) in N (A) and HPH (B). The percentage of circRNAs (y axis) was calculated according to the number of exons (x axis) spanned by each circRNA for the input circRNAs (left), m^6^A-circRNAs (red, right) and non-m^6^A circRNAs (blue, right) in N (C) and HPH (D). Up to seven exons are shown.

### The distribution and functional analysis for host genes of circRNAs with differentially expressed (DE) m^6^A peaks

The length of DE m^6^A circRNAs was mostly enriched in 1-10000 bps (Fig 3A). The host genes of upregulated m^6^A circRNAs were located in chromosome 1, 2 and 10, while the downregulated parts were mostly located in chromosome 1, 2 and 14 (Fig 3B).

**Fig 3.**
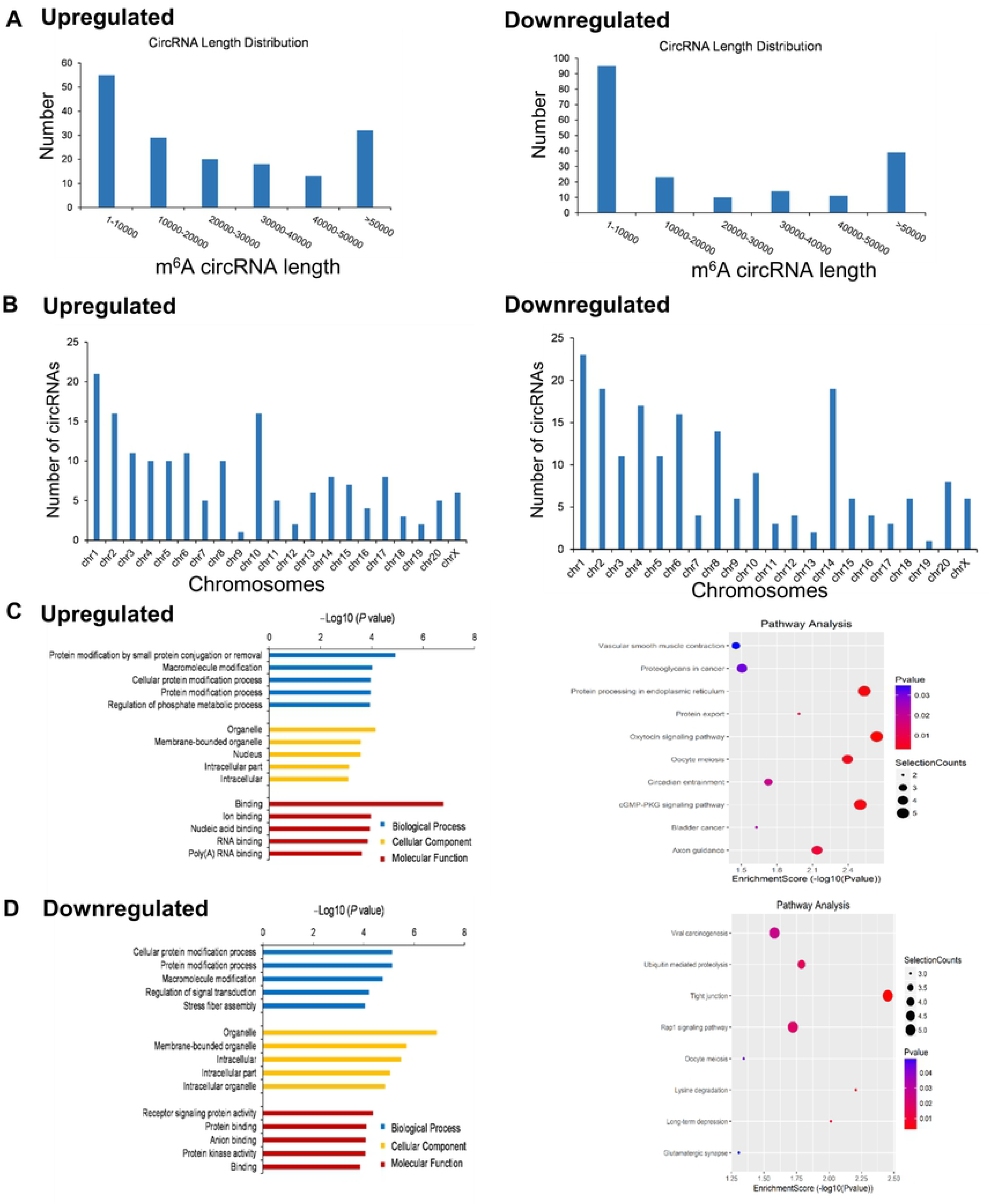
The distribution and functional analysis for host genes of circRNAs with differentially expressed (DE) m^6^A peaks. (A) DE m^6^A circRNAs length. (B) The chromosomes origins for host genes of DE m^6^A circRNAs. GO enrichment and KEGG signaling pathway analysis for host genes of upregulated (C) and downregulated (D) m^6^A circRNAs. GO enrichment analysis include biological process (BP) analysis, cellular component (CC) analysis, and molecular function (MF) analysis. P values are calculated by DAVID tool.

Gene ontology (GO) analysis and Kyoto Encyclopedia of Genes and Genomes (KEGG) pathway analysis were performed to explore the host genes of circRNAs with DE m^6^A peaks. In the GO analysis (Fig 3C, **left**), the parent genes of circRNAs with upregulated m^6^A peaks were enriched in the protein modification by small protein conjugation or removal and macromolecule modification process in the biological process (BP). Organelle and membrane-bounded organelle were also the two largest parts in the cellular component (CC) analysis. Binding and ion binding were the two main molecular functions (MF). The top 10 pathways from KEGG pathway analysis were selected in the bubble chart (Fig 3C, **right**). Among them, the oxytocin signaling pathway, protein processing in endoplasmic reticulum and cGMP-PKG signaling pathway were the top 3 pathways involved. In addition, vascular smooth muscle contraction pathway was the most associated pathway in PH progression (27).

In Fig 3D **left**, the parent genes of circRNAs with downregulated m^6^A peaks were mainly enriched in the cellular protein modification process and protein modification process in BP. Organelle and membrane-bounded organelle made up the largest proportion in the CC classification. The MF analysis was focused on receptor signaling protein activity and protein binding. The parent genes of circRNAs with decreased m^6^A peaks were mainly involved in the tight junction and lysine degradation in the KEGG pathway analysis (Fig 3D, **right**).

### m^6^A level of circRNAs and circRNAs abundance were influenced by hypoxia

360 m^6^A circRNAs were detected in N and HPH groups. 49% of circRNAs were only modified by m^6^A in N, and 54% of circRNAs were only modified by m^6^A in HPH (Fig 4A). To explore whether m^6^A methylation would influence circRNAs expression level, expression of the 360 common m^6^A circRNAs were identified. More circRNAs tended to decrease in HPH compared to N (Fig 4B). Moreover, expression of m^6^A circRNAs was significantly downregulated compared with non-m^6^A circRNAs in hypoxia, suggesting that m^6^A may downregulate the expression of circRNAs in hypoxia (Fig 4C, **p = 0.0465**).

**Fig 4.**
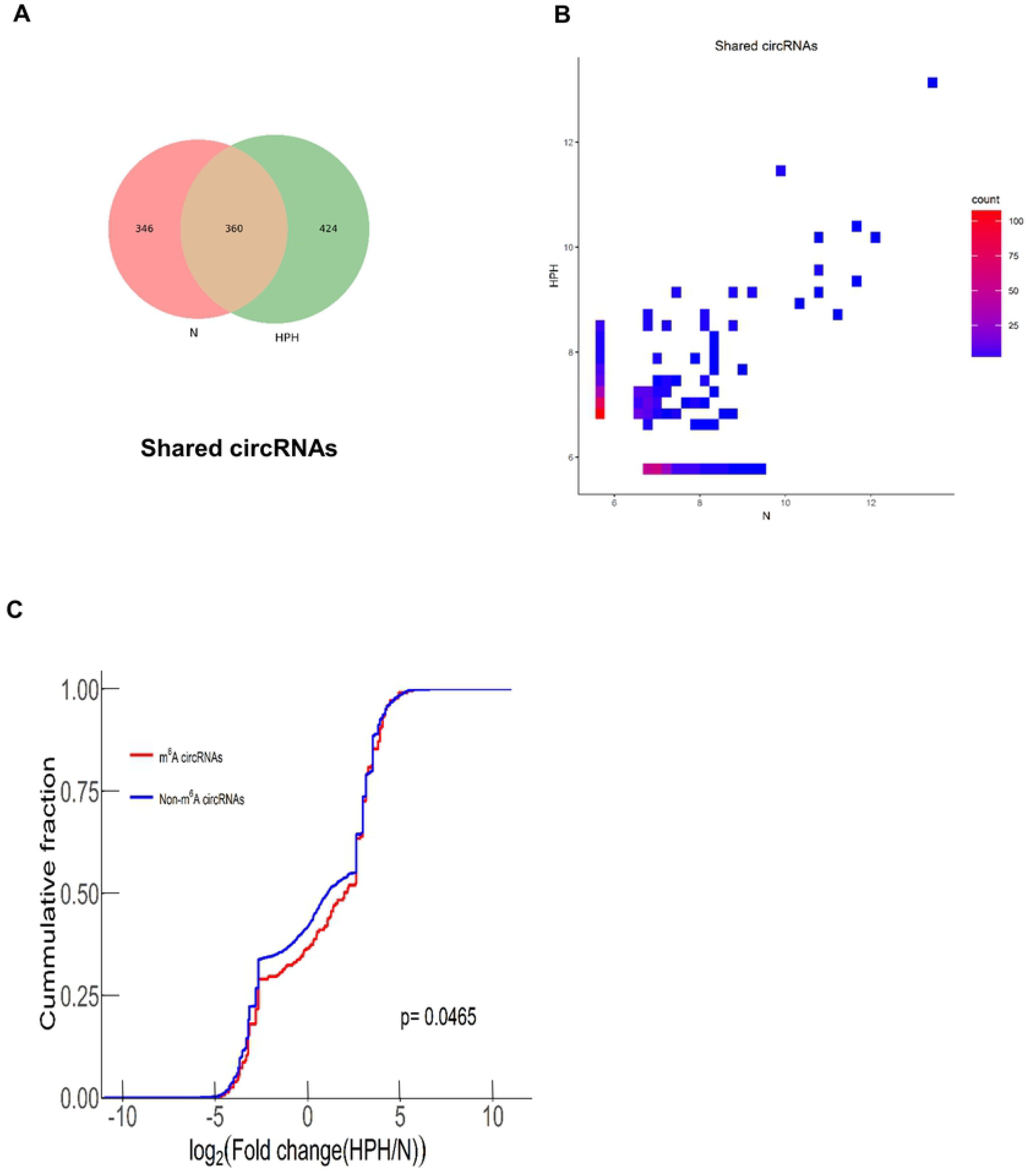
The relationship of m^6^A level of circRNAs and circRNAs abundance in hypoxia. (A) Venn diagram depicting the overlap of m^6^A circRNAs between N and HPH. (B) Two-dimensional histograms comparing the expression of m^6^A circRNAs in lungs of N and HPH rats. It showed that m^6^A circRNAs levels for all shared circRNAs in both groups. CircRNAs counts were indicated on the scale to the right. (C) Cumulative distribution of circRNAs expression between N and HPH for m^6^A circRNAs (red) and non-m^6^A circRNAs (blue). P value was calculated using two-sided Wilcoxon-Mann-Whiteney test.

### Construction of a circRNA–miRNA–mRNA co-expression network in HPH

We found 76 upregulated circRNAs with increased m^6^A abundance, and 107 downregulated circRNAs with decreased m^6^A abundance (Fig 5A, **S2 Table**). As known, circRNAs were mostly regarded as a sponge for miRNAs and regulated the expression of corresponding target genes of miRNAs (28). To explore whether circRNAs with DE m^6^A abundance influence the availability of miRNAs to target genes, we selected DE circRNAs with increased or decreased m^6^A abundance. GO enrichment analysis and KEGG pathway analysis were also performed to analyze target mRNAs. Target mRNAs displayed similar GO enrichment in the two groups (Fig 5B and 5C). Two main functions were determined in BP analysis: positive regulation of biological process and localization. Intracellular and intracellular parts make up the largest proportion in CC part. Target mRNAs were mostly involved in protein binding and binding in MF part. In the KEGG pathway analysis, the top 10 most enriched pathways were selected (Fig 5D and 5E). Wnt and FoxO signaling pathways were reported to be involved in PH progression (29–31). Then, we analyzed the target genes involved in these two pathways (**S1 Fig and S2 Fig**). SMAD4 was associated with PH and involved in Wnt signaling pathways. MAPK3, SMAD4, TGFBR1, and CDKN1B were involved in FoxO signaling pathways. To explore the influence of circRNA-miRNA regulation on PH-associated genes expression, we constructed a circRNA-miRNA-mRNA network, integrating matched expression profiles of circRNAs, miRNAs and mRNAs (Fig 5F and 5G). MicroRNAs sponged by the target genes of interest were analyzed. MiR-125a-3p, miR-23a-5p, miR-98-5p, let-7b-5p, let-7a-5p, let-7g-5p, and miR-205 were analyzed because they were reported to be associated with PH (32, 33). We filtered the key mRNAs and miRNAs, and founded that the two circRNAs were the most enriched, which were originated from chr1:204520403-204533534- (Xpo6) and chr7:40223440-40237400- (Tmtc3).

**Fig 5.**
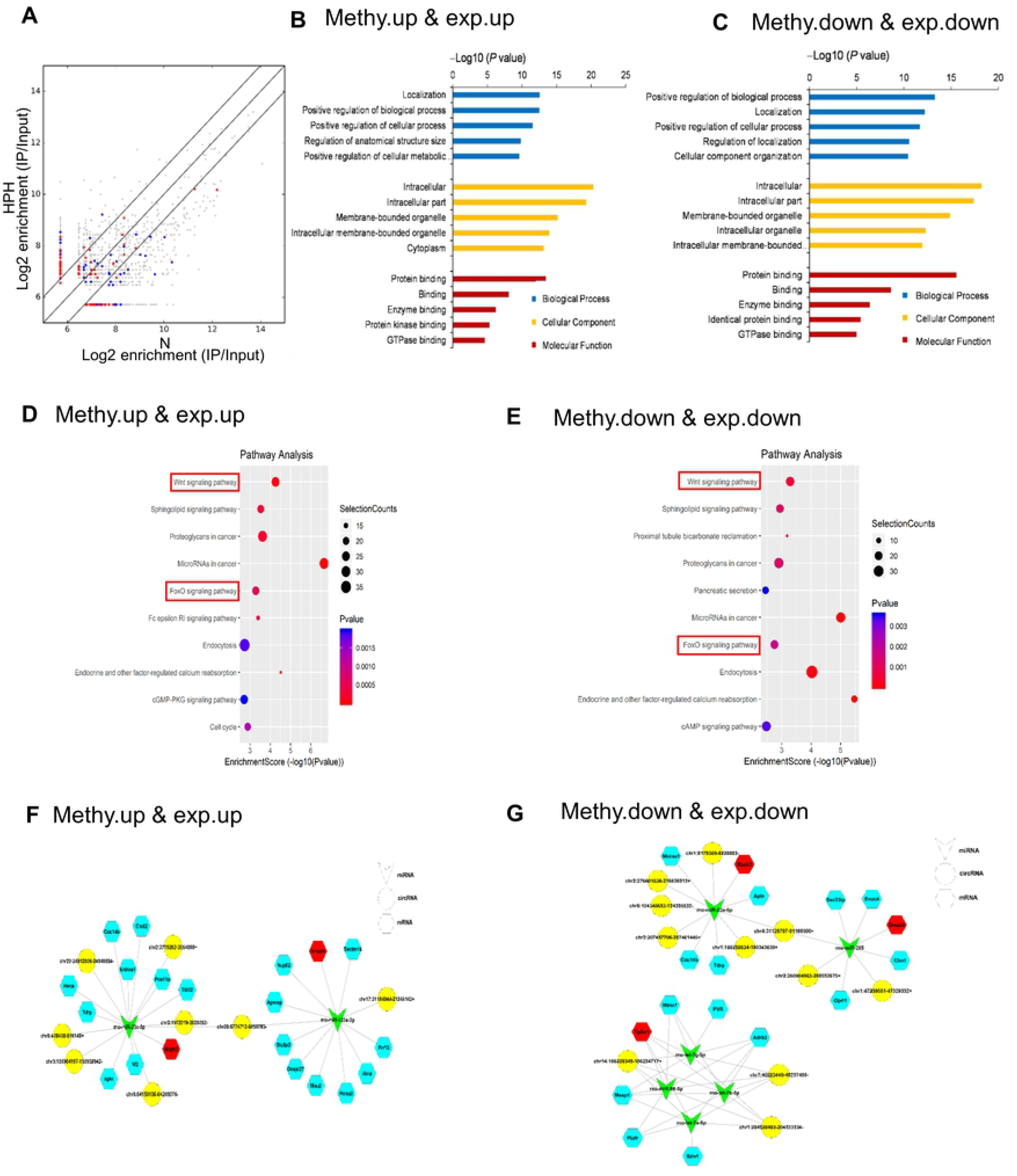
Construction of a circRNA–miRNA–mRNA co-expression network in HPH. (A) Comparison of the relationship between m^6^A level and expression of circRNAs between N and HPH. The fold-change ≥2.0 was considered to be significant, which was the abundance of m^6^A peaks of HPH relative to N. Red dots represents circRNAs with upregulated m^6^A level and blue dots represents circRNAs with downregulated m^6^A level. IP/Input referred to the abundance of m^6^A peak in circRNAs detected in MeRIP-Seq (IP) normalized to that detected in input. (B and C) GO enrichment analysis includes BP analysis, CC analysis, and MF analysis. P values are calculated by DAVID tool. (D and E) KEGG signaling pathway analysis for the downstream mRNAs which was predicted to be ceRNA of DE cirRNAs. Methy. down & exp. down represents downregulated cirRNAs with decreased m^6^A level. Methy. up & exp. up represents upregulated cirRNAs with increased m^6^A level. (F and G) CeRNA analysis for DE circRNAs. Network map of circRNA-miRNA-mRNA interactions. Green V type node: miRNA; yellow circular node: DE circRNAs; blue hexagon node: target genes of miRNAs; red hexagon node: PH-related genes.

### m^6^A circXpo6 and m^6^A circTmtc3 were downregulated in PASMCs and PAECs in hypoxia

m^6^A abundance was significantly reduced in PASMCs and PAECs when exposed to hypoxia (0.107% ± 0.007 vs. 0.054% ± 0.118, p = 0.023 in PASMCs; 0.114% ± 0.011 vs. 0.059% ± 0.008, p = 0.031 in PAECs, Fig 6A). m^6^A abundance in circRNAs was lower than it in mRNAs (0.1–0.4%) (17, 18). Next, we confirmed the back-splicing of circXpo6 and circTmtc3 by CIRI software. The sequence of linear Xpo6 and Tmtc3 mRNA was analyzed. Then we identified that circXpo6 was spliced form exon 7, 8, and 9 of Xpo6. CircTmtc3 was spliced form exon 8, 9, 10, and 11 (Fig 6B). Using cDNA and genomic DNA (gDNA) from PASMCs and PAECs as templates, circXpo6 and circTmtc3 were only amplified by divergent primers in cDNA, while no product was detected in gDNA (Fig 6C). To identify whether circXpo6 and circTmtc3 were modified by m^6^A, we performed m^6^A RNA Immunoprecipitation (MeRIP)-RT-PCR and MeRIP quantitative RT-PCR (MeRIP-qRT-PCR) to detect the expression of circXpo6 and circTmtc3 (Fig 6D and 6E). m^6^A circXpo6 and m^6^A circTmtc3 were significantly decreased in PASMCs and PAECs when exposed to hypoxia (p = 0.002, and p = 0.015 in PASMCs and p = 0.02, and p = 0.047 in PAECs)

**Fig 6.**
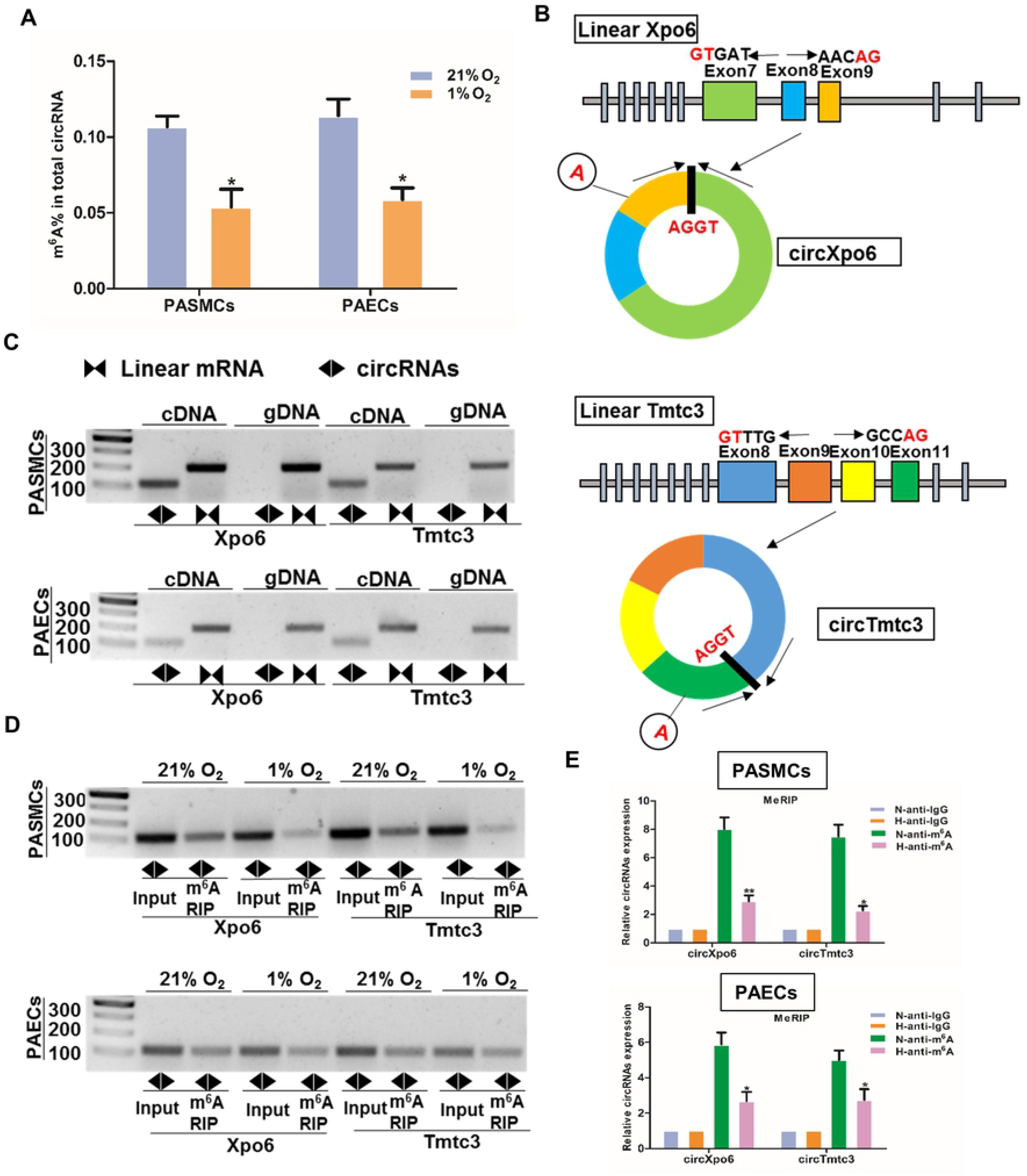
The expression profiling of m^6^A circXpo6 and m^6^A circTmtc3 in PASMCs and PAECs in hypoxia. (A) Box-plot for m^6^A peaks enrichment in circRNAs *in vitro*. Pulmonary arterial smooth muscle cells (PASMCs) and pulmonary artery endothelial cells (PAECs) were exposed to 21% O_2_ and 1% O_2_ for 48 h. Total RNA was extracted and treated by RNase R. m^6^A levels were determined as a percentage of total circRNAs. (B) Schematic representation of exons of the Xpo6 and Tmtc3 circularization forming circXpo6 and circTmtc3 (black arrow). (C) RT-PCR validation of circXpo6 and circTmtc3 in PASMCs and PAECs exposed to 21% O_2._ Divergent primers amplified circRNAs in cDNA, but not in genomic DNA (gDNA). The size of the DNA marker is indicated on the left of the gel. (D and E) RT-PCR and qRT-PCR was performed after m^6^A RIP in PASMCs and PAECs exposed to 21% (N) and 1% O_2_ (H) for 48 h. Input was used as a control (D). IgG was used as a negative control (E). Values are presented as means ± SD. *p ≤ 0.05 (different from 21% O_2_ or the N-anti-m^6^A); **0.001 ≤ p ≤ 0.009 (different from the N-anti-m^6^A).

## Discussion

In this study, we identified the transcriptome-wide map of m^6^A circRNAs in hypoxic pulmonary hypertension. On the whole, we found that m^6^A level in circRNAs was reduced in lungs when exposed to hypoxia. m^6^A circRNAs were mainly derived from single exons of protein-coding genes in N and HPH. m^6^A abundance in circRNAs was downregulated in hypoxia *in vitro*. m^6^A influenced the circRNA–miRNA–mRNA co-expression network in hypoxia. Moreover, circXpo6 and circTmtc3 were the novel identified circRNAs modified by m^6^A in hypoxic pulmonary hypertension.

m^6^A plays important roles in various biological processes. m^6^A is associated with cancer progression, promoting the proliferation of cancer cells and contributing to the cancer stem cell self-renewal (18, 21). Lipid accumulation was reduced in hepatic cells when m^6^A abundance in peroxisome proliferator-activator (*PPaR)* was decreased (34). Enhanced m^6^A level of mRNA contributed to compensated cardiac hypertrophy (35). Also, m^6^A modification of lincRNA 1281 was necessary for mESC differentiation (36).

Although it has been reported that m^6^A mRNAs were influenced by hypoxia, there is no report about m^6^A circRNAs in HPH yet. Up to now, no consistent conclusion was reached about the link between m^6^A and hypoxia. Previous reports found that the m^6^A abundance in mRNA was increased under hypoxia stress in HEK293T cells and cardiomyocytes (37, 38). The increased m^6^A level stabilized the mRNAs of Glucose Transporter 1 (Glut1), Myc proto-oncogene bHLH transcription factor (Myc), Dual Specificity Protein Phosphatase 1 (Dusp1), Hairy and Enhancer of Split 1 (Hes1), and Jun Proto-Oncogene AP-1 Transcription Factor Subunit (Jun) without influencing their protein level (37). In contrast, another reported that m^6^A level of total mRNA was decreased when human breast cancer cell lines were exposed to 1% O_2_ (26). Hypoxia increased demethylation by stimulating hypoxia-inducible factor (HIF)-1α- and HIF-2α–dependent over-expression of ALKBH5 (26). In addition, transcription factor EB activates the transcription of ALKBH5 and downregulates the stability of METTL3 mRNA in hypoxia/reoxygenation (H/R)-induced autophagy in ischemic diseases (38). Our study found that m^6^A abundance in total circRNAs was decreased by hypoxia exposure. Moreover, our study indicated that circXpo6 and circTmtc3 were the novel identified circRNAs modified by m^6^A in HPH. m^6^A abundance in circXpo6 and circTmtc3 was decreased in hypoxia. It is probably because of HIF-dependent and ALKBH5-mediated m^6^A demethylation (26).

Previous reports indicated that m^6^A methylation close to 3’UTR and stop codon of mRNA is inversely correlated with gene expression (14, 39). Low m^6^A level is negatively associated with circRNAs expression, while high m^6^A level is not linked to circRNAs expression in hESCs and HeLa cells (14). Consistent with the previous reports (14, 39), our study found that m^6^A reduced the total circRNAs abundance in hypoxia. Surprisingly, the expression of circXpo6 and circTmtc3 was decreased with the downregulated m^6^A level. No associated reports could confirm this phenomenon yet. Therefore, we suspected that m^6^A may influence the expression of circXpo6 and circTmtc3 through other pathways. It needs further validation.

Competing endogenous RNA (ceRNA) mechanism was proposed that mRNAs, pseudogenes, lncRNAs and circRNAs interact with each other by competitive binding to miRNA response elements (MREs) (40, 41). m^6^A acts as a post-transcript regulation of circRNAs and influences the expression of circRNAs, thus we suggested that m^6^A could also regulate the circRNA–miRNA–mRNA co-expression network. When the circRNAs were classified, we found that these downstream targets regulated by circRNA–miRNA of interest were mostly enriched in PH-associated Wnt and FoxO signaling pathways (30, 31). The Wnt/β-catenin (bC) pathway and Wnt/ planar cell polarity (PCP) pathway are the two most critical Wnt signaling pathways in PH (30). As known, the two important cells associated with HPH are PASMCs and PAECs (1, 3). The growth of PASMCs was increased when Wnt/bC and Wnt/PCP pathways were activated by platelet derived growth factor beta polypeptide b (PDGF-BB) (30, 42). In addition, the proliferation of PAECs was enhanced when Wnt/bC and Wnt/PCP pathways were activated by bone morphogenetic protein 2 (BMP2). Furthermore, the FoxO signaling pathway is associated with the apoptosis-resistant and hyperproliferative phenotype of PASMCs (31). Reactive oxygen species (ROS) is increased by hypoxia and activates AMPK-dependent regulation of FoxO1 expression, resulting in increased expression of catalase in PASMCs (43). Our study firstly uncovered that m^6^A influenced the stability of circRNAs, thus affecting the binding of circRNAs and miRNA, resulting in the activation of Wnt and FoxO signaling pathways.

However, limitations still exist in the study. First, we did not analyze the m^6^A level between circRNAs and the host genes. Second, the exact mechanism of hypoxia influences m^6^A was not demonstrated. Lastly, the function of circXpo6 and circTmtc3 in HPH was not elaborated.

In conclusion, our study firstly identified the transcriptome-wide map of m^6^A circRNAs in HPH. m^6^A level in circRNAs was decreased in lungs of HPH and in PASMCs and PAECs exposed to hypoxia. m^6^A level influenced circRNA–miRNA–mRNA co-expression network in HPH. Moreover, we firstly identified two downregulated m^6^A circRNAs in HPH: circXpo6 and circTmtc3. We suggest that circRNAs can be used as biomarkers because it is differentially enriched in specific cell types or tissues and not easily degraded (6). Also, the aberrant m^6^A methylation may contribute to tumor formation and m^6^A RNAs may be a potential therapy target for tumor (17). Therefore, we suppose that m^6^A circRNAs may also be used as a potential diagnostic marker or therapy target in HPH in the future. But more research is needed to validate this possibility.

## Materials and Methods

### Hypoxia-induced PH rat model

Sprague-Dawley rats (SPF, male, 180-200 g, 4 weeks) were obtained from the Animal Experimental Center of Zhejiang University, China. Rats were maintained in a normobaric normoxia (FiO_2_ 21%) or hypoxic chamber (FiO_2_ 10%) for 3 weeks (3, 44). Rats were then isoflurane-anesthetized and sacrificed. Lung and heart tissues were removed and immediately frozen at liquid nitrogen or fixed in 4% buffered paraformaldehyde solution. All experimental procedures were conducted in line with the principles approved by the Institutional Animal Care and Use Committee of Zhejiang University.

### RVSP and RVH

RVSP was measured as below. Rats were isoflurane-anesthetized and right ventricle catheterization was performed through the right jugular using a pressure-volume loop catheter (Millar) as the previous reports (44–46). The ratio of [RV/ (LV + S)] was used as an index of RVH.

### Histological analysis

Lung tissues were embedded in paraffin, sectioned at 4 μm and stained with hematoxylin and eosin (H&E) and α-smooth muscle actin (α-SMA, 1:100, ab124964, Abcam, USA). The ratio of pulmonary small artery wall thickness and muscularization were calculated (3).

### Isolation and hypoxia-treatment of PASMCs and PAECs

PASMCs and PAECs were isolated using the methods according to previous reports (32, 47, 48). PASMCs and PAECs were cultured in Dulbecco’s modified Eagle’s medium (DMEM) supplemented with 10% fetal bovine serum (FBS) and 20% FBS for 48h, respectively (32, 49). The cells were incubated in a 37°C, 21% O_2_ or 1% O_2_–5% CO_2_ humidified incubator. PASMCs at 70–80% confluence in 4 to 7 passages were used in experiments. PAECs at 80–90% confluence in 4 to 5 passages were used in experiments (50).

### RNA isolation and RNA-seq analysis of circRNAs

Total RNA (10 mg) was obtained using TRIzol reagent (Invitrogen, Carlsbad, CA, USA) from lungs (1 g) of control and HPH rats. The extracted RNAs were purified with Rnase R (RNR07250, Epicentre) digestion to remove linear transcripts. Paired-end reads were harvested from Illumina Hiseq Sequence after quality filtering. The reads were aligned to the reference genome (UCSC RN5) with STAR software. CircRNAs were detected and annotated with CIRI software(51). Raw junction reads were normalized to per million number of reads (RPM) mapped to the genome with log2 scaled.

### MeRIP and Library Preparation

Total RNA was extracted as the methods described above. Then, rRNA was depleted following DNase I treatment. RNase R treatment (5 units/mg) was performed in duplicate with 5 mg of rRNA-depleted RNA input. Fragmented RNA was incubated with anti-m^6^A polyclonal antibody (Synaptic Systems, 202003) in IPP buffer for 2 hours at 4°C. The mixture was then incubated with protein A/G magnetic beads (88802, Thermo Fisher) at 4°C for an additional 2 hours. Then, bound RNA was eluted from the beads with N^6^-methyladenosine (PR3732, BERRY & ASSOCIATES) in IPP buffer and extracted with Trizol reagent (15596026, Thermo Fisher). NEBNext® Ultra™ RNA Library Prep Kit (E7530L, NEB) was used to construct RNA-seq library from immunoprecipitated RNA and input RNA. The m^6^A-IP and input samples were subjected to 150 bp paired-end sequencing on Illumina HiSeq sequencer. Methylated sites on circRNAs were identified by MetPeak software.

### Construction of circRNA–miRNA–mRNA co-expression network

The circRNA–miRNA–mRNA co-expression network was based on the ceRNA theory that circRNA and mRNA shared the same MREs. Cytoscape was used to visualize the circRNA–miRNA–mRNA interactions based on the RNA-seq data. The circRNA-miRNA interaction and miRNA–mRNA interaction of interest were predicted by TargetScan and miRanda.

### Measurement of Total m^6^A, MeRIP-RT-PCR and MeRIP-qRT-PCR

Total m^6^A content was measured in 200 ng aliquots of total RNA extracted from PASMCs and PAECs exposed to 21% O_2_ and 1% O_2_ for 48 h using an m^6^A RNA methylation quantification kit (P-9005, Epigentek) according to the manufacturer’s instructions. MeRIP (17-701, Millipore) was performed according to the manufacturer’s instruction. A 1.5 g aliquot of anti-m^6^A antibody (ABE572, Millipore) or anti-IgG (PP64B, Millipore) was conjugated to protein A/G magnetic beads overnight at 4^◦^C. A 100 ng aliquot of total RNA was then incubated with the antibody in IP buffer supplemented with RNase inhibitor and protease inhibitor. The RNA complexes were isolated through phenol-chloroform extraction (P1025, Solarbio) and analyzed via RT-PCR or qRT-PCR assays. Primers sequences were listed as follows: circXpo6, 5’ TCTGGGAGACAAGGAAGCAG3’ (forward) and 5’ CAGGATGGGGATGGGCTG3’ (reverse); circTmtc3, 5’ TACCCATGTTCAGCCAGGTT3’ (forward) and 5’ GAAGCCAAGCATTCACAGGA3’ (reverse); linear Xpo6, 5’ CTGTGTTTTGGGTCAGGAGC 3’ (forward) and 5’ ATCGAGTTCCTCTAGCCTGC3’ (reverse); linear Tmtc3, 5’ ACTCTGCTGTGATTGGACCA3’ (forward) and 5’ AGAAGAGGTTTGATGCGGGA3’ (reverse).

### Data analysis

3’ adaptor-trimming and low quality reads were removed by cutadapt software (v1.9.3). Differentially methylated sites were identified by the R MeTDiff package. The read alignments on genome could be visualized using the tool IGV. DE circRNAs were identified by Student’s *t*-test. GO and KEGG pathway enrichment analysis were performed for the corresponding parental mRNAs of the DE circRNAs. GO enrichment analysis was performed using the R topGO package. KEGG pathway enrichment analysis was performed according to a previous report (52). GO analysis included BP analysis, CC analysis, and MF analysis. MicroRNAs sponged by the target genes were predicted by TargetScan and microRNA. P values are calculated by DAVID tool for GO and KEGG pathway analysis. The rest statistical analyses were performed with SPSS 19.0 (Chicago, IL, USA) and GraphPad Prism 5 software (La Jolla, CA). N refers to number of samples in figure legends. The statistical significance was determined by Student’s *t*-test (two-tailed) or two-sided Wilcoxon-Mann-Whiteney test. P < 0.05 was considered statistically significant. All experiments were independently repeated at least three times.

## Acknowledgements

We thanked all subjects who participated in this study.

## Supporting information

**S1 Fig. KEGG pathway analysis for Wnt and FoxO signaling pathway in methy. up & exp. up group.**

**S2 Fig. KEGG pathway analysis for Wnt and FoxO signaling pathway in methy. down & exp. down group.**

**S1 Table. Differentially expressed m^6^A abundance in circRNAs.**

**S2 Table. Differentially expressed m^6^A abundance linked with differentially expressed circRNAs abundance.**

